# Molecular characterization of clonal human renal forming cells

**DOI:** 10.1101/2020.03.05.978254

**Authors:** Cohen-Zontag Osnat, Gershon Rotem, Harari-Steinberg Orit, Kanter Itamar, Omer Dorit, Pleniceanu Oren, Tam Gal, Oriel Sarit, Ben-Hur Herzl, Katz Guy, Zohar Dotan, Kalisky Tomer, Dekel Benjamin, Pode-Shakked Naomi

## Abstract

The adult kidney replaces its parenchyma *in vivo* in steady state and during regeneration by segment-specific clonal cell proliferation.

To understand human adult kidney clonal cell growth, we derived tissue from human nephrectomies and performed limiting dilution to establish genuine clonal cultures from one single cell.

Clonal efficiency of the human kidney was x%. Remarkably, a single renal cell could give rise to up to 3.3*10(6) cells. Phenotypically, two types of clonal cultures were apparent; a stably proliferating cuboidal epithelial-like appearing (EL) and a rapidly proliferating fibroblast-like appearing (FL). RNA sequencing of all clonal cultures separated FL from EL cultures according to proximal-distal/collecting renal epithelial tubular identity, respectively. Moreover, distinct molecular features in respect to cell-cycle, epithelial-mesenchymal transition, oxidative phosphorylation, BMP signaling pathway and cell surface markers were observed for each clone type. Surprisingly, clonal expansion (>3 months) was sustained in EL clones harboring markers of mature kidney epithelia (high CD24, CDH1, EpCAM, EMA) in contrast to de-differentiated FL clones (high NCAM1, serpine1), which showed fast lineage amplification and exhausted in a few weeks.

Thus, the human adult kidney harbors progenitor cell function in which segment identity and the level of epithelial differentiation dictate clonal characteristics.

## Introduction

With end stage renal disease becoming a worldwide challenge and subsequently the need for kidney donors expanding, far exceeding their availability, there is a burning need for alternative regenerative approaches. For both tissue engineering and direct cellular interventions, utilization of cell sources harboring bona a fide renal potential, e.g. renal lineage cells that function to generate or replace mature kidney cells, may prove beneficial [1].

In the developing kidney there exists a pool of self-renewing mesenchymal multipotent nephron progenitors that give rise to various nephron epithelial cell types to generate whole nephrons [2, 3]. Human nephrogenesis ceases at 34 weeks of gestation as we are born with a fixed number of nephrons and are unable to initiate whole nephrogenesis in adulthood [4]. Previously, we obtained a comprehensive view of *in vivo* mouse kidney epithelial cell dynamics at the single-cell level [5]. Using multicolored fluorescent fate-mapping mouse models to track the behavior of single epithelial cells and clonal units, we discovered that cell turn-over in the nephron relies on multiple foci of clonal proliferation of kidney-resident epithelial cells that are dispersed along the nephron. Long-term *in vivo* analysis of clonal progeny showed each clone to contribute to a single renal epithelial lineage indicating that adult kidney growth was segmented and that kidney resident epithelial cells act as unipotent progenitors for local regeneration [4]. Thus, clonal proliferation is a mechanism of cell replacement in the adult kidney. Similarly, using lineage tracing in mice, LGR5+ cells were shown to mark distal tubular epithelial progenitors in development [6], while Schutgens et al. showed that cells expressing the stem cell marker *Tnfrsf19* (also known as *Troy*), clonally contribute to tubular structures during development and continue to give rise to segment-specific collecting duct cells in homeostasis of the adult kidney [7].

Analyzing kidney repair following acute damage, intrinsic proximal tubule cell proliferation was shown to possess a major role in replacing the lost cell mass [8, 9]. In addition, several evidence support epithelial dedifferentiation to ensue kidney injury [10-12]. KIM1/HAVCR1 is a specific marker for injured proximal tubule cells [11]. Recently, in vivo lineage tracing of proximal tubule KIM1+ cells following acute kidney injury, revealed co-expression of KIM1, VIMENTIN, SOX9, and KI67 in their derived clones, suggesting concomitant mesenchymal dedifferentiated and proliferative states of these cells [8].

Alternatively, some evidence support a specific subset of ‘professional’ tubular progenitors (for instance PAX2 expressing cells) to also contribute to tubular repair after acute injury [13].

Nevertheless, the principles governing genuinely single cell derived clonal adult kidney growth ex vivo have hardly been dealt with. Herein we developed a culturing system to enable clonal proliferation and propagation of single kidney epithelial cells derived from human adult kidneys coupled with transcriptional characterization. In contrast to bulk culture growth which quickly deteriorates following epithelial-mesenchymal transition (EMT), single cell clones could be maintained for over 3 months in culture. Our results disclose distinct types of proximal and distal single-cell kidney derived clones harboring specific molecular characteristics that correspond to *in vivo* tubular cell traits and unveil possible molecular drivers and markers for human kidney clonal cell growth.

## Materials and Methods

### Ethics Statement

This study was conducted according to the principles expressed in the Declaration of Helsinki. The study was approved by the Institutional Review Boards of Sheba Medical Center, Wolfson Hospital and Asaf Harofeh Medical Center. All procedures were carried out following signed informed consents.

All pregnant women involved in the study provided written informed consent for the collection of samples from their aborted fetuses and subsequent analysis.

### Establishment of primary cultures from the human kidney

Human adult kidney samples were recovered from healthy margins of renal cell carcinoma (RCC) tumors, resected from partial or total nephrectomies [1, 14]. Human Fetal Kidneys (hFKs) at gestational ages ranging from 15 to 22 weeks were harvested from fetuses that were electively aborted as previously described [15]. Collected tissues were washed with cold HBSS (Invitrogen) and minced into ∼1 mm cubes using sterile surgical scalpels followed by incubation at 37°C for two hours in Iscoves’ Mod Dulbecco’s Medium (IMDM) (Invitrogen) supplemented with 0.1% collagenase IV (Invitrogen). The processed tissue was then forced through 100 µm strainers to achieve a single cell suspension. After the digesting medium was removed, the cells were resuspended in a growth medium and plated in flasks. Serum-free medium (SFM) was comprised of N2 medium (Biological Industries) supplemented with 1% Pen-strep 100M, 1% L-glutamine, 0.4% B27 supplement (Gibco), 4µg/ml heparin sodium (Intramed), 1% non-essential amino acids, 1% sodium pyruvate, 0.2% CD Lipid concentrate (all from Invitrogen), 2.4mg/ml glucose, 0.4mg/ml transferrin, 10mg/ml insulin, 38.66µg/ml putrescine, 0.04% sodium selentine, 12.6µg/ml progesterone (all from Sigma-Aldrich), 10ng/ml FGF and 20ng/ml EGF. For passaging, cell detachment was performed via non-enzymatic cell dissociation solution (Sigma-Aldrich). Cells were observed and photographed using Nikon Eclipse TS100 and Nikon Digital Sight camera (Figure S1).

### Preparation of Conditioned Medium

Conditioned medium (CM) was prepared by cultivating hFK cells in SFM for 48-72h, using an inoculum density of 3.0 × 10^5^ cells mL^−1^. Spent medium was harvested by centrifugation, and the supernatant sterile filtered through a 0.22 μm bottle-top filter (Lab design) and kept at −20 °C until used.

### Single-cell cloning by limiting dilution assay and self-renewal assay

Cells were detached as described above and diluted with culturing medium. Cells were then plated onto pre-coated 96 well plates with Matrigel (BD Biosciences) at a density of 0.3 or 1 cell per with either 100% SFM or in 50% conditioned medium (CM) and 50% SFM. After 3-4 weeks the number of colonies was evaluated for each dilution. Clusters of cells were considered colonies when they were visible macroscopically. Colonie forming efficiency (CFE) was determined from the formula CFE (%) = (number of colonies/number of cells seeded) × 100. Upon confluence, single cell clones were detached using cell dissociation solution and re-plated in a larger fibronectin coated plate (Sigma-Aldrich) with SFM.

### Assessment of cell growth

4,000 cells were plated in triplicates and grown in 96-well plates over-night. The following day the medium was changed and supplemented with various concentrations of GW9662 (Sigma-Aldrich) or vehicle only (DMSO) as control, and the cells were further incubated for 48 or 96 hours. Cell proliferation was measured using CellTiter 96 Aqueous One Solution Cell Proliferation Assay (Promega) according to the manufacturer’s instructions. Briefly, the cells were incubated with the MTS solution for 3 hours at 37°C and absorbance at 492 nm was determined using the infinite F50 microplate reader (Tecan). Three independent experiments were carried out for each source.

### Doubling time and growth rate calculations

Approximately 80,000 cells were seeded in 6-well plates. The following day and every day thereafter, cells were trypsinized and counted using a hemocytometer. Calculation of doubling time and growth rate (defined as the number of cell divisions per time unit) was carried out using a verified online calculator [16] quantifying the cells’ doubling time, based on the number of cells in at least 3 time points.

### RNA sequencing and analysis

Bulk total RNA was quantified on an Agilent BioAnalyzer (Agilent Technologies) and aliquots of 270-500ng were made into cDNA libraries using the TruSeq mRNA-Seq library kit (Illumina). Libraries were then sequenced 1 × 50 bases on the Illumina HiSeq 2500 platform.

Sequence data was analyzed using the protocol by Anders et al [17]. Briefly, raw reads were aligned by TopHat2 [18] to the human hg19 genome. Aligned reads were counted by HTSeq [19]. Data normalization and differential gene expression was done by DESeq2 [20]. Gene enrichment analysis was performed with GSEA [21]. Differentially expressed genes with P-value < 0.05 and absolute log2 fold change >0.5 were considered for downstream analysis. Euclidian distance and PCA analysis were performed on 1000 most variable genes. We used R packages pheatmap, and ggplot2 to generate the heatmaps, and all other plots in this study.

### Statistical analysis

Error bars represent the mean ± SD, unless otherwise indicated. Statistical differences between cell populations were evaluated using the non-parametric, one sided sign test. Statistical differences between two group data were analyzed via Student’s t test. For all statistical analysis, the level of significance was set as p<0.05.

## Results

### Phenotypical and functional characterization of single cell-derived colonies from the human adult kidney (hAK)

Human adult kidney cells derived from human nephrectomies were used for establishment of single cell clones. Cells were seeded in 96 well plates and cultured at a density of 0.3 cells per well [10] and grown in defined SFM medium for 2 to 4 weeks. Single cell clones were initially characterized according to their size as large (L) medium (M) or small (S) clones. Colony Forming Efficiency (CFE)% was 10.29±4.95 globally, S%=4.69, M%=2.47, L%=3.125 (Figure 1A). We next assessed in vitro self-renewal and expansion potential via serial passaging of the generated clones. Interestingly, most colonies surviving passaging and expansion were large at passage (Pc) 0 (100% at Pc1 and 33.33% that continued to expand for up to 4 passages) compared to medium colonies that presented limited ability for clonal expansion (63% at Pc1, X% at Pc2 and none that continued to expand over to Pc3) and small colonies that failed to survive along passages altogether (Figure 1B-C). Accordingly, large colonies that survived were highly proliferative [as manifested by their lower doubling time (DT)] in contrast to the heterogeneous pool of cultured cells from which the colonies were derived, which were already undergoing senescence at the same time point (Figure 1D). Colonies originating from a single cultured AK cell demonstrated the potential to expand into approximately 3.27E+06 cells (Figure 1E). Clones maintained their original morphology along passages for more than 3 months (Figure 1F). A second subdivision of the single cell epithelial clones was done according to their morphology. Two distinct morphologic subgroups were observed; either cuboidal epithelial-like (EL) or spindle shaped fibroblast-like (FL) clones (Figure 1G). Most EL clones (5 out of 6) were large and four were able to propagate in culture as opposed to only one large propagatable FL clone out of three. Thus, single cell derived hAK clones can be phenotypically characterized according to clone size and morphology. Propagatable clones preserved their original phenotype during passages (see also supplementary figure S1).

**Figure 1.**
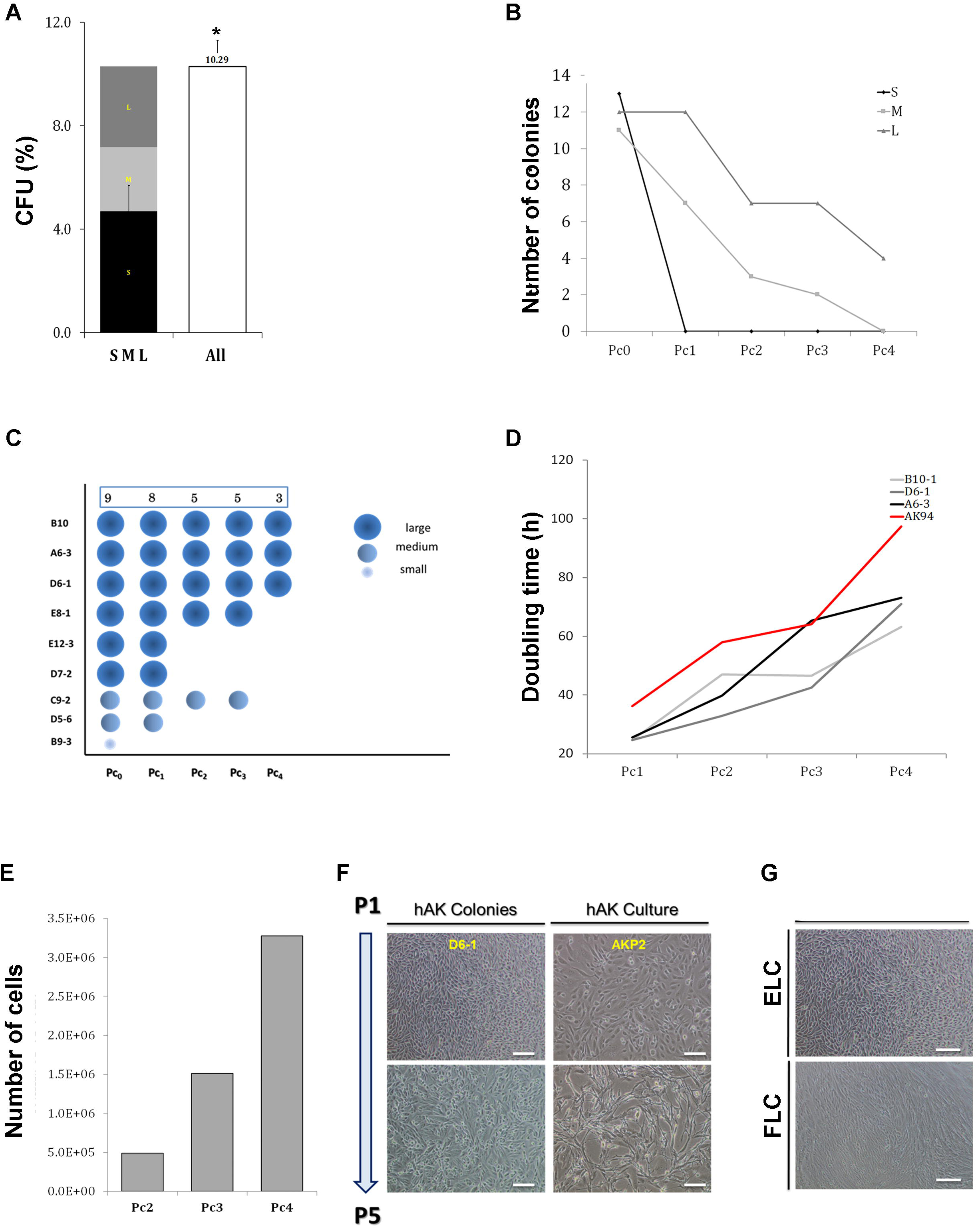
Establishment and characterization of single cell-derived colonies from human kidney. (A) CFE of single cell clones derived from fresh hAK cells according to their size (CFE%=10.29±4.95); *P<0.01; Small (S)%=4.69, Medium (M)%=2.47, Large (L)%=3.125; (B) Representative graph of self-renewal capacity of hAK-clones base on colony size. Data are presented for each passage as relative number of clones generated from the total number of cells plated. Most colonies surviving expansion were large (L) colonies (100% at Pc1 and 33.33% continued to expand for up to 4 passages) compared to medium (M) colonies that presented limited ability for clonal expansion (63% at Pc1 and none continued to expand beyond the 3^rd^ passage) and small (S) colonies that failed to survive along passages altogether; (C) Representative scheme of 9 hAK colonies over passages according to size. Each dot represents a specific clone and the size of the dot represents clone size. The numbers above indicate the survived clone in each passage. As can be seen, clones surviving expansion were large colonies (100% survived Pc1 and 33.33% continued to expand for up to 4 passages) compared to medium colonies that presented limited ability for clonal expansion (in hAK 63% at Pc1 and none that continued to expand for up to 4 passages) and small colonies that failed to survive along passages; (D) Representative graph of hAK large clones’ growth rate during expansion according to doubling time (DT). Large colonies that survived were highly proliferative compared to the heterogeneous pool of cultured cells from which they were derived; (E) Representative graph of the expansion potential of hAK single cell derived clones. Clones originating from a single cultured AK cell demonstrated were able to expand into approximately 3.27E+06 cells; (F) Representative morphology of an expanded hAK single cell derived clone (D6-1) from along passages, compared to the heterogeneous hAK culture from which the clones was derived. A stable epithelial (EL) phenotype was preserved during clonal expansion for several month in contrast to the heterogeneous pool of cultured cells, which were already undergoing senescence and switching to a fibroblast-like morphology at the same time point. Scale bars, 100μm; (G) Representative photomicrographs of single cell clonal phenotypes. Two type of clones were generated: Epithelial-like (ELC) or Fibroblast-like (FLC). Scale bars, 100 μm.

### RNA-Sequencing analysis defines clone types upon in vitro expansion of human Adult Kidney clones

In order to characterize hAK clones at the molecular level we have performed RNA-Sequencing of 9 hAK clones. Bulk hAK cultures were used as control. Initial examination of EL, FL clones and bulk hAK cultures showed that early bulk hAK cultures (early hAK) undergo major transcriptional changes during propagation into late cultures (late hAK), while clonal propagation, as reflected by the transition from early into late clones, results in significantly smaller overall transcriptional changes (Figure 2A). Moreover, FL clones clustered with late hAK cultures, while all EL clones clustered together, distinct from the former, supporting a phenotypic-transcriptomic correlation and the maintenance of a stable transcriptome concomitant with the observed stable phenotype of the EL clones (Figure 2B).

**Figure 2.**
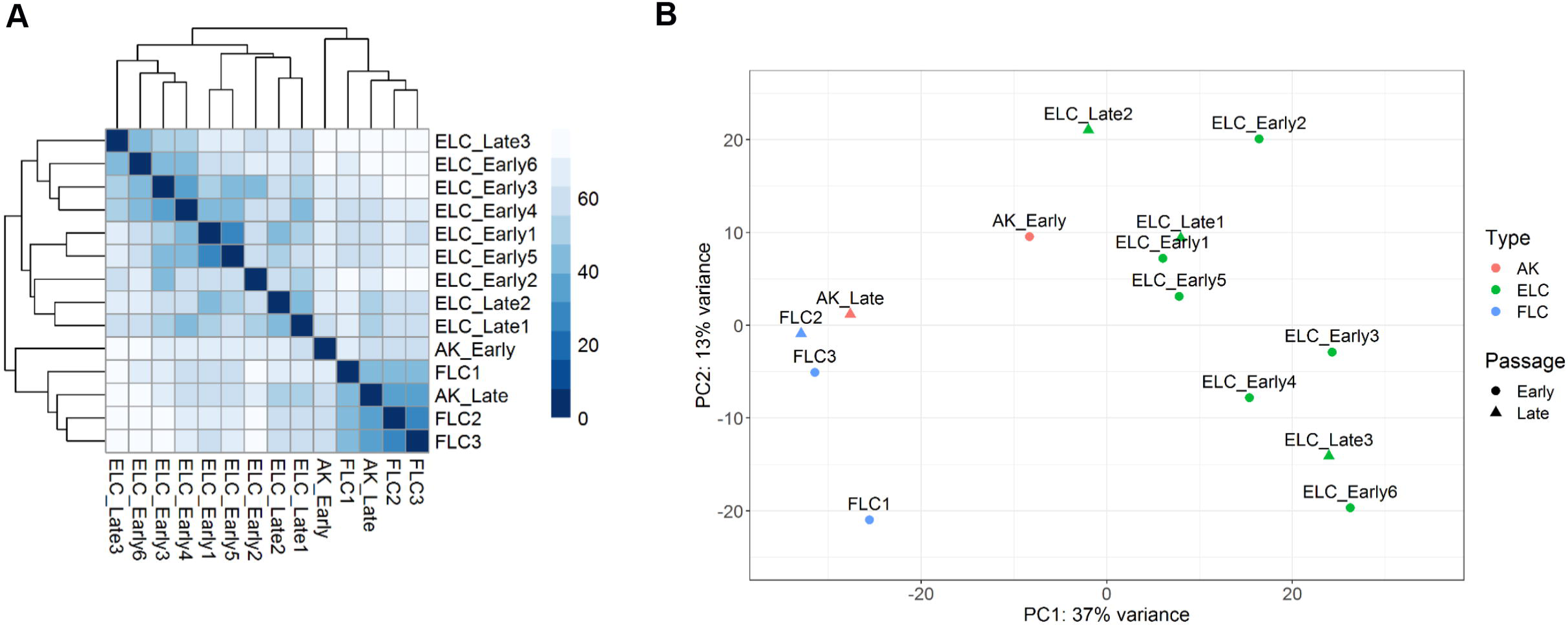
RNA-Sequencing analysis defines clone types upon in vitro expansion of human Adult Kidney single cell derived clones. (A) Euclidean distance analysis was performed on RNA-seq data to illustrate variation in transcription levels between samples. Variation in transcription levels varies greatly between early bulk human adult kidney (BAK) and late BAK, while early single cell clones show lower variation in transcriptional levels along in vitro propagation into late clones; FLCs are more similar to late BAK than to ELCs while early and late ELCs are clustered together. The distance between cell types is demonstrated by a heatmap. Darker color indicates lesser distance which means greater similarity; (B) Principal component analysis was performed on RNA-seq data to illustrate variation in transcription levels between samples. Different colors denote culture type. Different shapes denote early or late passage. Bulk Human Adult Kidney-BAK; FLC-Fibroblast-Like Clones; ELC-Epithelial-Like Clones;

### Molecular characterization of clone types reveal differences in the degree of epithelial maturation and nephron segment identity

In order to deepen our understanding of the driving forces preserving clonal epithelial growth, we sought to characterize the two morphologically distinct single cell clone types (i.e. EL and FL) at the molecular level. Therefore, we have analyzed genes that were differentially expresses between the two clone types. Analysis of tissue-specific genes, which mark different nephron segments revealed the EL clones to possess a more distal tubular expression signature as manifested by elevation of SLC12A3, EMA, MUC1 etc., while FL clones demonstrated a proximal tubular gene expression (i.e. high ENPEP, AQP1, CD13, WT1, HVCR1 etc.) (Figures 3A, B). While EL clones demonstrated preservation of epithelial gene expression including CDH1, GRHL2, EPCAM and KRT7 (Figure 3C, F), FL clones showed elevated EMT genes including NCAM1, FGF1, SERPINB2, FN1, ZEB2 as well as cell-cycle genes along with downregulation of CD24 (Figure 3C, D, F). Additionally, oxidative phosphorylation genes were significantly higher in FL compared with EL clones (Figure 3E, F). Concomitantly, gene ontology (GO) analysis revealed enrichment for mitochondrial related genes (i.e. electron transport, mitochondrial protein complex etc.) in line with mitochondrial abundancy of proximal tubular cells (Figure S2). Cluster of differentiation (CD) genes were differentially expressed between EL and FL clones in line with their previously described renal segment specificity and location along the EMT axis (Figure 3G). Intriguingly, several CD genes with unique characteristics were also differentially expressed between the two clone types. These included, NCAM (CD56) a marker of early mesenchymal stem cells in the embryonic kidney that was highly expressed by FL clones. Additionally, CD109, a GPI-anchored protein that plays an impotent role in TGF beta signaling [22], was up-regulated in FL clones. In contrast, immune response related CDs (i.e. CD46, CD55, CD274) were highly expressed by EL clones. Moreover, EL clones expressed high levels of CD9, an exosome-specific marker.

**Figure 3.**
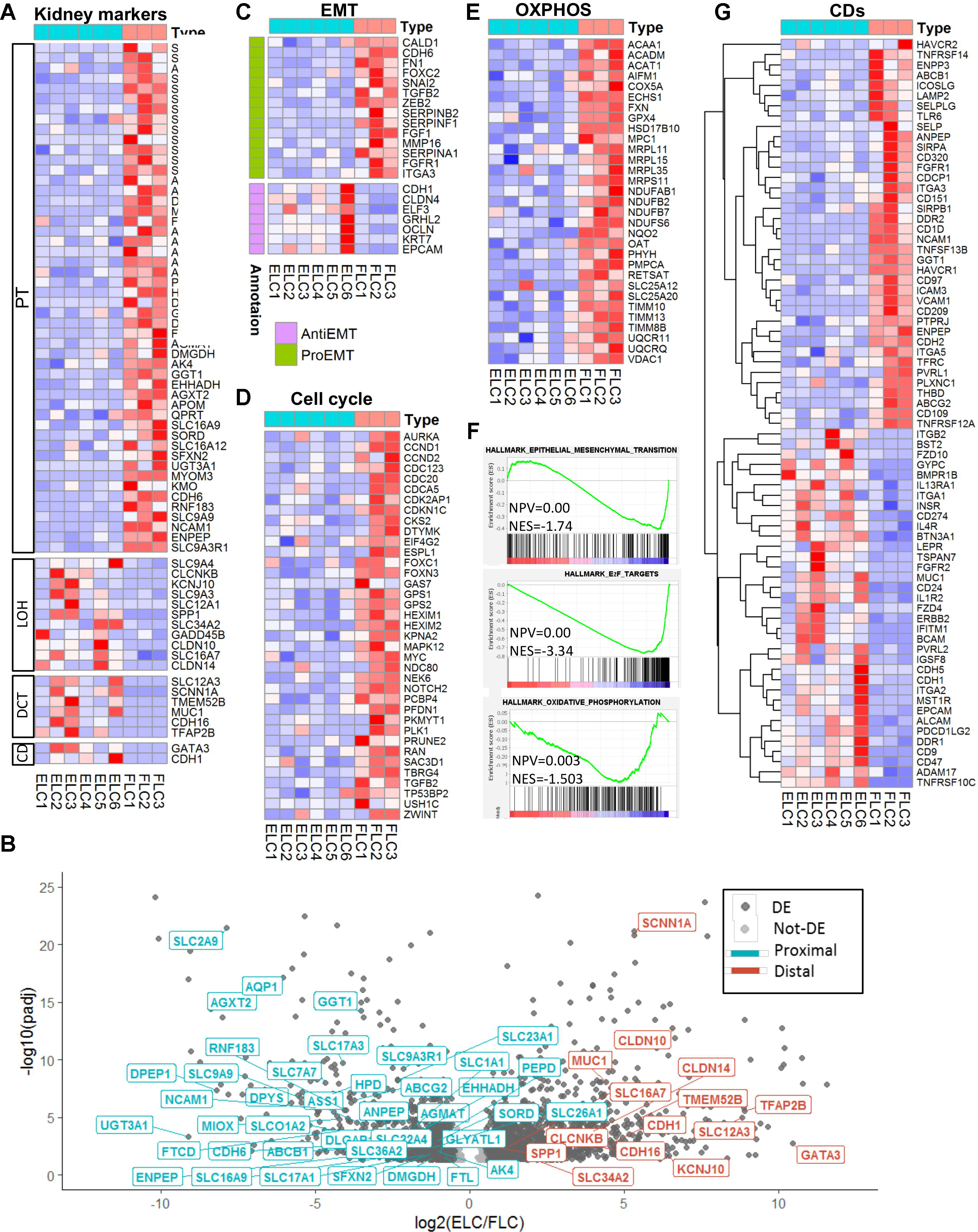
Molecular characterization of clone types reveal differences in the degree of epithelial maturation and nephron segment identity. HeatMap representation of differentially expressed human nephron segment specific markers between FLC and ELC. FLC demonstrate a proximal nephron while ELC show distal nephron gene signature (A); Volcano plot representation of genes deferentially expressed in ELC in comparison to FLC. Dark-grey points denote differentially expressed genes (padj<0.05, Log2(fold-change)>0.5). Kidney specific genes from different segments of the nephrons are labeled. While genes that mark the proximal segment of the nephron are down-regulated in ELC (cyan), genes that mark more distal segments of the nephron are up-regulated in ELC (red) (B); HeatMap representation of differentially expressed EMT (C), cell-cycle (D) and oxidative phosphorylation (E) genes between FLC and ELC, showing upregulation in FLC compared to ELC; Gene set enrichment analysis (GSEA) showing enrichment of cell-cycle, EMT and oxidative phosphorylation genes in FLC (F); HeatMap representation of differentially expressed CD genes between FLC and ELC (G); Abbreviations: ELC- Epithelial-like clones; FLC– Fibroblast-like clones; EMT- epithelial to mesenchymal transition; CD- cluster of differentiation.

As EL clones demonstrated preferable in vitro propagation capacity while preserving their phenotype for over 3 months, we next looked for possible “drivers” for clonal propagation. The most dominant signal transduction pathway in EL clones was the BMP/SMAD pathway, previously shown to take play a significant role in tubular regeneration as well as prevent fibrotic signaling in renal tubular cells following kidney injury [22]. Elevation in BMP2, BMP4, BMP6, SMAD7, SMAD6, among other members of this pathway, was observed in EL clones (Figure S3), putting it forward as a potential driver for human kidney epithelial clonal expansion. Thus, FL clones show EMT, cell-cycle, embryonic stem cell and oxidative phosphorylation genes while EL clones maintain epithelial maturation markers and activate BMP signaling pathway.

Renal developmental genes may be activated in adult renal cells [23], therefore we queried transcription factors (TFs) important for nephrogenesis in our transcriptome data. EL clones showed expression of TFs expressing in ureteric bud (GATA3, HOXB7, HOXB9, GRHL2) and also mesenchyme (MM) TFs (HOXD10, SALL2, PBX1) (Figure S4A). On the contrary, FL clones disclose MM TFs (WT1, HNF1A, FOXC1) (Figure S4A). Thus, EL clones may originate from both MM and UB, suggesting that distal tubule clones maybe overlapping with those of collecting system clones.

SLC17A3 and VCAM1 are proximal markers found to be upregulated in clear cell renal cell carcinoma (ccRCC), both are highly elevated in FL clones (Figure S4B). Interestingly, ccRCC markers such as ITGA5, GGT1, CDH6 and CA9 were also found to be upregulated in FL clones (Figure S4B). ccRCC is also characterized by EMT, cell cycle activation and oxidative phosphorylation [24-26] further supporting the resemblance between ccRCC and FL clones.

### During propagation EL-clones acquire FL-clonal transcriptional signature while preserving distal tubular segment identity

In order to dissect the transcriptional processes taking place in EL clones during passages we also analyzed the transcriptome of late EL clones grown in culture for three months.

While late EL clones revealed preservation of renal distal-segment identity (Figure 4A), a transcriptomic shift towards FL clonal signature was observed. This included an elevation in cell cycle genes (i.e. CCND1. CCND2, MYC, etc.) (Figure 4B) and Epithelial to Mesenchymal transition (EMT) genes (i.e. FGF1, SERPINF1, SERPINB2, ZEB2 etc.) (Figure 4C). Intriguingly, even though a shift towards EMT was demonstrated in late EL clones, several distinct epithelial markers were preserved during EL clonal propagation including CDH1, EPCAM, MUC1 etc. (Figure 4C). During in vitro propagation, EL clones display marked elevation of oxidative phosphorylation genes is in parallel to that observed in FL clones (Figure 4D). Closer look at the expression of CD genes during in vitro propagation revealed preserved expression of CD9, CD46, CD24 among others, while CD274, CDFZD4, FZD10, BMPR1 genes were downregulated and CD151, CD97, CD320 genes, were upregulated, resembling the expression pattern seen in FL clones (Figure 4E).

**Figure 4.**
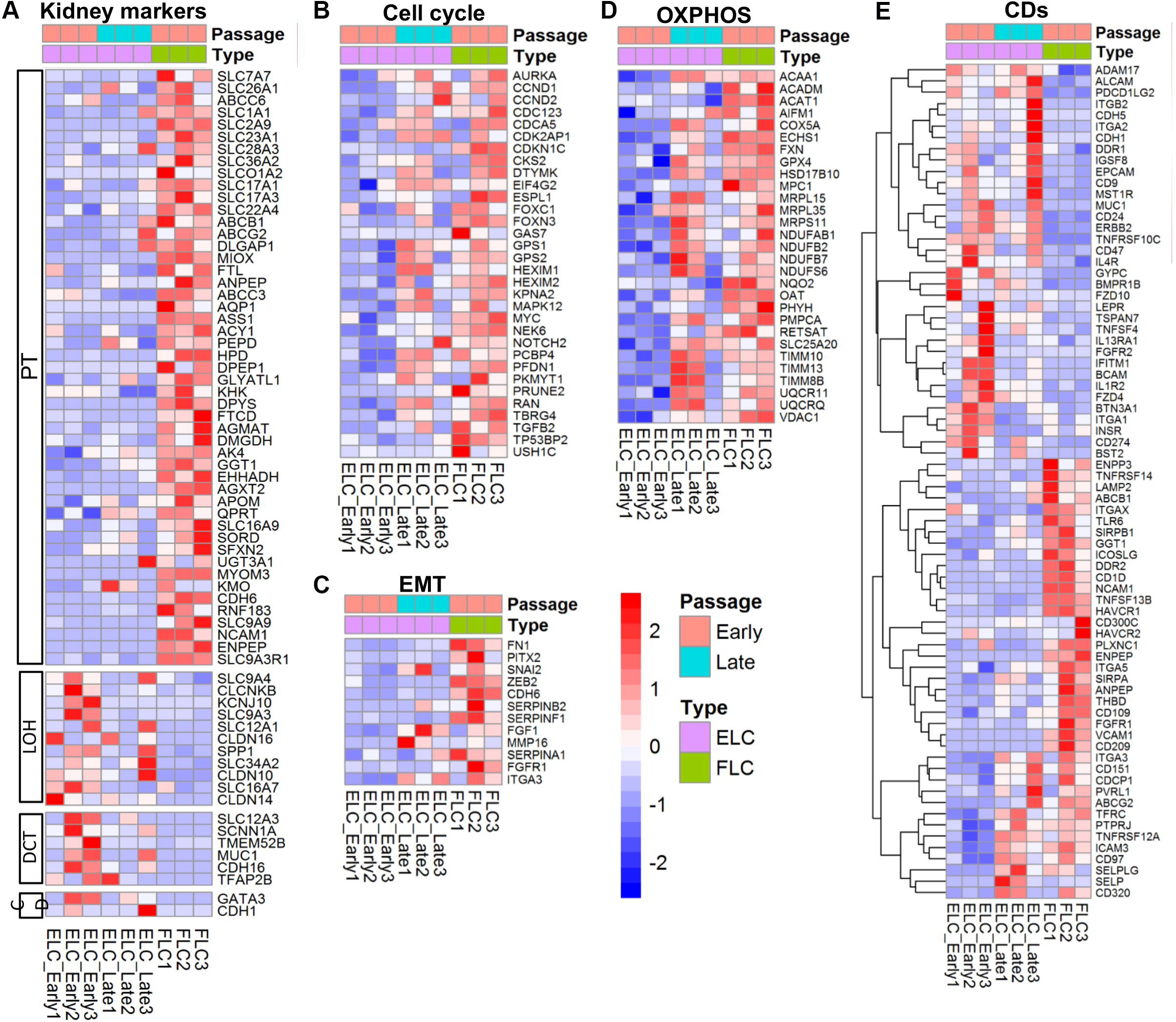
During propagation, EL-clones acquire FL-clonal transcriptional signature while preserving distal nephron segment identity. HeatMap representations of differentially expressed genes between early ELC, late ELC and FLC. Along in vitro expansion, ELC demonstrate preservation of renal distal-segment identity (A); transcriptomic shift towards FL clonal signature in cell-cycle (B), EMT (C), and oxidative phosphorylation (D) genes; Cluster of differentiation (CD) genes show preservation of CD9, CD46, CD24 among others alongside downregulation of CD274, CDFZD4, FZD10, BMPR1 and upregulation of CD151, CD97, CD320, during in vitro expansion of ELC clones (E). Abbreviations: ELC- Epithelial-like clones; FLC– Fibroblast-like clones; EMT- epithelial to mesenchymal transition;

In conclusion, during in vitro propagation, EL clones preserve their original renal distal segment identity along with several epithelial markers, while concomitantly acquiring FL clonal transcriptional signature.

## Discussion

Previous studies have highlighted *in vivo* clonal behavior of tubular cells as a major characteristic for cell replacement in the adult kidney [22]. Herein, we have calibrated a novel and reliable culture system that allows *ex-vivo* single cell derived clonal growth of the human kidney in order to delineate its phenotypic and molecular characteristics.

We demonstrate that some clones not only proliferate, but also maintain a stable phenotype and quiescence-proliferation balance as manifested by constant replication time, over consecutive passages. This point is not at all trivial, considering the significant de-differentiation, degeneration and senescence undergone by parallel heterogeneous hAK culture in the same period, proving the unique potential of the clonal assay. Remarkably, clonal human kidney cultures harbor stem/progenitor characteristics enabling clonal growth over 3 months and clonal output of up to 3.3*10(6) differentiated renal cells that eventually far exceeds the *in vivo* mouse system.

Phenotypically two distinct epithelial single cell derived clonal cultures were observed. The first showed fibroblastic appearance, initial proliferation burst, generating clones of variable size, with a reduced propagation capacity over passages. The second demonstrate cuboidal epithelial characteristics, have a steady proliferation rate, usually forming larger clones and are able to propagate over several passages in culture. This clonal heterogeneity was resolved by RNA sequencing of single cell derived clonal cultures; All FL type clones represent clones originating from the proximal tubules as manifested by high expression of proximal tubule markers such as ENPEP, AQP1, and CD13 among others (see Figure 3). In contrast, EL clones derive from distal renal segments displaying overexpression of distal tubule markers (i.e. SLC12A3, MUC1, CDH1 among others). This striking separation ignited in depth molecular analysis of clone types. Concomitant with a proximal tubule transcriptional signature, FL clones demonstrated enhanced EMT, cell cycle and embryonic stem cell gene expression, suggestive of concomitant proliferative and de-differentiation states. Interestingly, the regenerative response to acute kidney injury is dependent on proximal tubular cells undergoing concurrent dedifferentiation and proliferation [7, 8]. Thus, transcriptional signature of FL clones mimics the regenerative response of proximal tubular cells to an acute kidney insult. In this regard, FL clones may inform on the clonal *in vivo* regenerative response; For instance, while Ncam1 (CD56) has been previously indicated as a marker activated in a subset of cultured proximal tubular cells with clonogenic properties [3, 27] and here indeed shown to be a molecular marker of proximal clonal proliferation. CD109 and CD209 are noteworthy to mention both representing migratory markers that are upregulated in cancer [28, 29]. Specifically, CD109 is a member of the TGF beta signal transduction, negatively regulating TGF beta/SMAD complex in keratinocytes [30]. It was recently put forward as a marker for cancer stem-like cells in epithelioid sarcoma and as a poor prognostic marker in soft tissue sarcomas [31]. Epithelioid sarcoma are rare malignant tumors of mesenchymal origin that histologically display mix epithelial and mesenchymal traits [32]. Taken together, expression of these CDs might explain the fibroblast-like, mesenchymal characteristics of FL clones derived from a single kidney epithelial cell. Recent single cell RNA-seq of clear cell RCC disclosed a proximal tubular genetic signature with specifically elevated SLC17A3 and VCAM1. These molecules were overexpressed in FL clones along with ITGA5, GGT1, CDH6 and CA9 also previously shown to be highly expressed by clear cell RCC [33-37]. Thus, intriguingly proximal clonal proliferation shares markers of clear cell RCC. This may suggest that proximal clonal proliferation is a precursor cell lesion for RCC.

Oxidative phosphorylation, respiratory chain, mitochondrial matrix and envelope genes were also significantly elevated in these clones. Thus, clones do not appear to switch to aerobic glycolysis (Warburg effect [38]) as to be expected for highly proliferation cells, but rather increase oxidative phosphorylation. This observation is in line with inherent mitochondrial abundancy of proximal tubule cells, relying on aerobic respiration in order to be able to account for their high metabolic demands (reabsorption of 80% of the glomerular filtrate), whereas distal parts of the nephron possess a greater capacity to shift from oxidative phosphorylation to glycolysis [11]. In addition, increased oxidative phosphorylation was recently put forward as harboring a significant role in cell proliferation through a process of mitochondrial fusion [39].

In contrast, distal tubule transcriptional characteristics of the EL clones were accompanied by high epithelial maturation markers (i.e. CDH1, EPCAM, KRT7, GRLH2 etc.) alongside relatively decreased expression of EMT and cell cycle genes. Since more quiescent EL clones possess increased in vitro propagation capacity compared to FL clones, it seems that the degree of epithelial differentiation is crucial for clonal propagation. This contradicts the model whereby stem/progenitor function is linked to un-differentiated cells lacking maturation markers [40] and points towards stem/progenitor behavior as being a functional state [40]. High CD24 expression coincides with epithelial maturation markers in EL clones and as such may represent a marker associated with epithelial-type clonal proliferation. Interestingly, Elevation of CD genes related to immune response and exosomal communication was noted in EL clones. The former included CD46 and CD55, associated with compliment pathway retardation [41] and CD274, linked with inhibition of T cell function, all three were shown to decrease kidney allograft rejection [42, 43]. CD9, an exosome-complex gene was also elevated in EL clones. It was shown to take part in exosomal communication between the proximal and distal tubular segments in the human kidney, resulting in decreased production of reactive oxygen species (ROS) in distal tubular cells [44]. Thus, alongside distal tubular cells being inherently more resistant to ischemia as well as to oxidative stress [45], this CD expression pattern suggest a possible protective role during epithelial-type clonal growth.

Signal transduction pathways are selectively activated in different parts of the nephron during development and in response to renal insult. During EL clonal growth, BMP signaling was the most prominent pathway activated (i.e. high BMP2, BMP4, BMP6, SMAD1, SMAD6 expression) (Figures 3 and S3).

BMP signaling is involved in nephron patterning during kidney development and later on, operates selectively along specific nephron segments [46]. For instance, BMP4 is essential for mesoderm formation and patterning [46, 47]. Recently, BMP4 was shown to contribute to branching of the collecting duct in kidney organoid formation [48]. In accordance with this pathway activation in EL clones that maintain both epithelial phenotype and epithelial transcriptional signature, several BMP proteins (i.e. BMP2, BMP4, BMP6, BMP7 etc.) were shown to possess an anti-fibrotic effect following acute kidney injury [49-51]. Moreover, activation of the BMP-SMAD pathway is seen in epithelial cells that are re-lining the nephron following acute injury [50].

Additionally, in a model of interstitial nephritis, induced inactivation of the ALK3 receptor (a high-affinity receptor for BMP2 and BMP4) sensitizes proximal tubule cells to injury and promotes fibrosis [50].

Taken together, activation of the BMP signaling pathway in the unique pattern observed in EL clones corresponds to its role in kidney development as well as its anti-fibrotic effects. On the other hand, downregulation of this pathway in FL clones is in accordance with the pro-fibrotic response of proximal tubule cells to injury in its absence.

Importantly, a consecutive analysis on transcriptional changes during EL clonal propagation revealed preservation of segment identity while concomitantly exhibiting a transcriptional shift towards the FL-clonal signature. These changes were manifested as elevation in MET and cell cycle genes alongside a metabolic shift towards oxidative phosphorylation and occurred irrespective of the preservation of a cellular appearing epithelial phenotype. Hence, it remains to be determined whether a continued BMP signal or potentially mitochondrial augmentation discovered by drug screens of clonal cultures analyzing clonal proliferation can inhibit diversion from an EL to FL clonal signature (or perhaps direct an early FL signature to an EL counterpart).

To conclude, we show that *in vitro* single cell growth of the human adult kidney can be utilized to phenocopy, at least in part, it’s in vivo characteristics. The ability to function as adult stem/progenitor cells, clonally proliferate at the single cell level and propagate is related to the cell’s epithelial differentiation status favoring distal over proximal identity. Moreover, the methodology described herein, enabling long term in vitro culturing of human adult kidney cells that maintain quiescence-proliferation balance and preserve renal identity, provides a unique source to study replicating human kidney cells.

## Acknowledgements

This work was supported by the Israel Science Foundation (grant no. 910/20111) and 2071/2017, NIH DIACOMP, Sheba Medical Center (grant no. 25732-64), Bretler Foundation, Sackler school of Medicine; I.K., S.O., and T.K., were supported by the Israel Science Foundation (ICORE no. 1902/12 and Grants no. 1634/13 and 2017/13), the Israel Cancer Association (Grant no. 20150911), the Israel Ministry of Health (Grant no. 3-10146), and the EU-FP7 (Marie Curie International Reintegration Grant no. 618592).

## Author Contributions

B.D., R.G., O.HS, D.O. and N.PS., designed the experiments; R.G., S.O., H.BH., Z.D. G.K. and N.PS. performed the experiments. O.CZ, T.K., O.HS., I.K., T.K., G.T. N.PS. and B.D. analyzed the data; N.PS, B.D., R.G., O.CZ, O.P. and T.K. wrote the manuscript.

---

**Figure S1 |** Schematic representation of the method used for establishment of an efficient and reliable method for forming single cell clones from primary human kidney. Flow diagram of the steps involved in adult kidney clones isolation and expansion.

**Figure S2** | (A) Gene set enrichment analysis of differentially expressed statistics comparing ELC to FLC reveal an enrichment of mitochondrial-related GO terms in genes down-regulated in ELC (blue) and an enrichment of GO-terms related to development and specification processes in genes up-regulated in ELC (red); (B) Schematic representation of changes in mitochondrial related genes in FLC and ELC showing most complexes to be highly expressed by FLC. Annotations: Abbreviations: ELC-Epithelial-like clones, FLC-Fibroblast-like clones.

**Figure S3** | Schematic representation of changes in BMP/SMAD signal transduction pathway genes, showing many of its members to be highly expressed in ELC. Abbreviations: ELC-Epithelial-like clones

**Figure S4|** (A) Expression levels of nephrogenesis transcription factors differentially expressed between Early EL clones (red) and FL clones (turquoise). While genes upregulated in ELC are expressed in both the UB and the MM during renal development, genes upregulated in FLC are known to be expressed only in the MM; (B) Expression levels of Renal Cell Carcinoma (RCC) markers differentially expressed between Early EL clones (red) and FL clones (turquoise). Genes upregulated in FLC are known to be expressed in RCC.

**Table S1 |** Full RNA-Seq Data row counts used for the expression analysis of the different cultures (AK, ELC, FLC) generated from Early or Late passages.

## References

1 Harari-Steinberg, O., et al., Ex Vivo Expanded 3D Human Kidney Spheres Engraft Long Term and Repair Chronic Renal Injury in Mice. Cell Rep, 2020. 30(3): p. 852–869 e4.

2 Herzlinger, D., et al., Metanephric mesenchyme contains multipotent stem cells whose fate is restricted after induction. Development, 1992. 114(3): p. 565–72.

3 Harari-Steinberg, O., et al., Identification of human nephron progenitors capable of generation of kidney structures and functional repair of chronic renal disease. EMBO Mol Med, 2013. 5(10): p. 1556–68.

4 Pleniceanu, O., O. Harari-Steinberg, and B. Dekel, Concise review: Kidney stem/progenitor cells: differentiate, sort out, or reprogram? Stem Cells, 2010. 28(9): p. 1649–60.

5 Rinkevich, Y., et al., In vivo clonal analysis reveals lineage-restricted progenitor characteristics in mammalian kidney development, maintenance, and regeneration. Cell Rep, 2014. 7(4): p. 1270–83.

6 Barker, N., et al., Lgr5(+ve) stem cells drive self-renewal in the stomach and build long-lived gastric units in vitro. Cell Stem Cell, 2010. 6(1): p. 25–36.

7 Schutgens, F., et al., Troy/TNFRSF19 marks epithelial progenitor cells during mouse kidney development that continue to contribute to turnover in adult kidney. Proc Natl Acad Sci U S A, 2017. 1 :(52)14p. E11190–E11198.

8 Chang-Panesso, M., et al., FOXM1 drives proximal tubule proliferation during repair from acute ischemic kidney injury. J Clin Invest, 2019. 129(12): p. 5501–5517.

9 Chevalier, R.L., The proximal tubule is the primary target of injury and progression of kidney disease: role of the glomerulotubular junction. Am J Physiol Renal Physiol, 2016. 311(1): p. F145–61.

10 Bonventre, J.V., Dedifferentiation and proliferation of surviving epithelial cells in acute renal failure. J Am Soc Nephrol, 2003. 14 Suppl 1: p. S55–61.

11 Kramann, R., T. Kusaba, and B.D. Humphreys, Who regenerates the kidney tubule? Nephrol Dial Transplant, 2015. 30(6): p. 903–10.

12 Witzgall, R., et al., Localization of proliferating cell nuclear antigen, vimentin, c-Fos, and clusterin in the postischemic kidney. Evidence for a heterogenous genetic response among nephron segments, and a large pool of mitotically active and dedifferentiated cells. J Clin Invest, 1994. 93(5): p. 2175–88.

13 Lazzeri, E., et al,. Endocycle-related tubular cell hypertrophy and progenitor proliferation recover renal function after acute kidney injury. Nat Commun, 2018. 9(1): p. 1344.

14 Buzhor, E., et al., Kidney spheroids recapitulate tubular organoids leading to enhanced tubulogenic potency of human kidney-derived cells. Tissue Eng Part A, 2011. 17(17-18): p. 2305–19.

15 Pode-Shakked, N., et al., Evidence of In Vitro Preservation of Human Nephrogenesis at the Single-Cell Level. Stem Cell Reports, 2017. 9(1): p. 279–291.

16 V,. V.R.R. 2006. p. <http://www.doubling-time.com/compute.php.<

17 Humphreys, B.D., et al., Intrinsic epithelial cells repair the kidney after injury. Cell Stem Cell, 2008. 2(3): p. 284–91.

18 Kim, D., et al., TopHat2: accurate alignment of transcriptomes in the presence of insertions, deletions and gene fusions. Genome Biol, 2013. 14(4): p. R36.

19 Anders, S., P.T. Pyl, and W. Huber, HTSeq--a Python framework to work with high-throughput sequencing data. Bioinformatics, 2015. 31(2): p. 166–9.

20 Love, M.I., W. Huber, and S. Anders, Moderated estimation of fold change and dispersion for RNA-seq data with DESeq2. Genome Biol, 2014. 15(12): p. 550.

21 Subramanian, A., et al., Gene set enrichment analysis: a knowledge-based approach for interpreting genome-wide expression profiles. Proc Natl Acad Sci U S A, 2005. 102(43): p. 15545–50.

22 Cuttle, L., et al., Bcl-X(L) translocation in renal tubular epithelial cells in vitro protects distal cells from oxidative stress. Kidney Int, 2001. 59(5): p. 1779–88.

23 Metsuyanim, S., et al., Expression of stem cell markers in the human fetal kidney. PLoS One, 2009. 4(8): p. e6709.

24 Liu, J., et al., Loss of SETD2 Induces a Metabolic Switch in Renal Cell Carcinoma Cell Lines toward Enhanced Oxidative Phosphorylation.J Proteome Res, 2019. 18(1): p. 331–340.

25 Landolt, L., et al., Clear Cell Renal Cell Carcinoma is linked to Epithelial-to-Mesenchymal Transition and to Fibrosis. Physiol Rep, 2017. 5(11.(

26 Sanchez, D.J. and M.C. Simon, Genetic and metabolic hallmarks of clear cell renal cell carcinoma. Biochim Biophys Acta Rev Cancer, 2018. 1870(1): p. 23–31.

27 Buzhor, E., et al., Cell-based therapy approaches: the hope for incurable diseases. Regen Med, 2014. 9(5): p. 649–72.

28 Chuang, C.H., et al., Molecular definition of a metastatic lung cancer state reveals a targetable CD109-Janus kinase-Stat axis. Nat Med, 2017. 23(3): p. 291–300.

29 Figel, A.M., et al., Human renal cell carcinoma induces a dendritic cell subset that uses T-cell crosstalk for tumor-permissive milieu alterations. Am J Pathol, 2011. 179(1): p. 436–51.

30 Finnson, K.W., et al., Identification of CD109 as part of the TGF-beta receptor system in human keratinocytes. FASEB J, 2006. 20(9): p. 1525–7.

31 Emori, M., et al., High expression of CD109 antigen regulates the phenotype of cancer stem-like cells/cancer-initiating cells in the novel epithelioid sarcoma cell line ESX and is related to poor prognosis of soft tissue sarcoma. PLoS One, 2013. 8(12): p. e84187.

32 Sobanko, J.F., L. Meijer, and T.P. Nigra, Epithelioid sarcoma: a review and update. J Clin Aesthet Dermatol, 2009. 2(5): p. 49–54.

33 Bansal, A., et al., Gamma-Glutamyltransferase 1 Promotes Clear Cell Renal Cell Carcinoma Initiation and Progression. Mol Cancer Res, 2019. 17(9 :(p. 1881–1892.

34 Young, M.D., et al., Single-cell transcriptomes from human kidneys reveal the cellular identity of renal tumors. Science, 2018. 361(6402): p. 594–599.

35 Paul, R., et al., Cadherin-6, a cell adhesion molecule specifically expressed in the proximal renal tubule and renal cell carcinoma. Cancer Res, 1997. 57(13): p. 2741–8.

36 Paul, R., et al., Cadherin-6: a new prognostic marker for renal cell carcinoma. J Urol, 2004. 171(1): p. 97–101.

37 Soyupak, B., et al., CA9 expression as a prognostic factor in renal clear cell carcinoma. Urol Int, 2005. 74(1): p. 68–73.

38 Liberti, M.V. and J.W. Locasale, The Warburg Effect: How Does it Benefit Cancer Cells? Trends Biochem Sci, 2016. 41(3): p. 211–218.

39 Kumari, S., et al., Hyperglycemia alters mitochondrial fission and fusion proteins in mice subjected to cerebral ischemia and reperfusion. Transl Stroke Res, 2012. 3(2): p. 296–304.

40 Post, Y. and H. Clevers, Defining Adult Stem Cell Function at Its Simplest: The Ability to Replace Lost Cells through Mitosis. Cell Stem Cell, 2019. 25(2): p. 174–183.

41 Cheng, L., et al., Complement regulatory proteins in kidneys of patients with anti-neutrophil cytoplasmic antibody (ANCA)-associated vasculitis. Clin Exp Immunol, 2018. 191(1): p. 116–124.

42 Starke, A., et al., Renal tubular PD-L1 (CD274) suppresses alloreactive human T-cell responses. Kidney Int, 2010. 78(1): p. 38–47.

43 Michielsen, L.A., et al., Association Between Promoter Polymorphisms in CD46 and CD59 in Kidney Donors and Transplant Outcome. Front Immunol, 2018. 9: p. 972.

44 Gildea, J.J., et al., Exosomal transfer from human renal proximal tubule cells to distal tubule and collecting duct cells. Clin Biochem, 2014. 47(15): p. 89–94.

45 Gobe, G.C. and D.W. Johnson, Distal tubular epithelial cells of the kidney: Potential support for proximal tubular cell survival after renal injury. Int J Biochem Cell Biol, 2007. 39(9): p. 1551–61.

46 Oxburgh, L., et al., Bone morphogenetic protein signaling in nephron progenitor cells. Pediatr Nephrol, 2014. 29(4): p. 531–6.

47 Winnier, G., et al., Bone morphogenetic protein-4 is required for mesoderm formation and patterning in the mouse. Genes Dev, 1995. 9(17): p. 2105–16.

48 Mills, C.G., et al., Asymmetric BMP4 signalling improves the realism of kidney organoids. Sci Rep, 2017. 7(1): p. 14824.

49 Xia, Y., et al., Dragon enhances BMP signaling and increases transepithelial resistance in kidney epithelial cells. J Am Soc Nephrol, 2010. 21(4): p. 666–77.

50 Larman, B.W., et al., Chordin-like 1 and twisted gastrulation 1 regulate BMP signaling following kidney injury. J Am Soc Nephrol, 2009. 20(5): p. 1020–31.

51 Vigolo, E., et al., Canonical BMP signaling in tubular cells mediates recovery after acute kidney injury. Kidney Int, 2019. 95(1 :(p. 108–122.

