## Supplemental information for "Molecular characterization of clonal human renal forming cells"

Hashomer, Israel

^6^Division of Pediatric Nephrology, Edmond and Lily Safra Children's Hospital,

Sheba Medical Center, Tel-Hashomer, Israel

^7^Sackler Faculty of Medicine, Tel-Aviv University, Tel-Aviv, Israel

^8^L.E.M. Laboratory of Early Detection, Nes Ziona, Israel

^9^Department of Obstetrics and Gynecology, Assaf Harofeh Medical Center, Tzrifin, Israel

^*^These first authors contributed equally to this work

^#^These senior authors contributed equally to this work

^^^Correspondence:

Benjamin Dekel MD, PhD 
Pediatric Stem Cell Research Institute
Edmond & Lily Safra Children's Hospital, 
Sheba Medical Center 

### Supplemental information – Table of content:

Supplemental figures and legends	3

Figure S1|	3

Figure S2|	4

Figure S3|	5

Figure S4|	6

legend for Supplemental tables	7

Table S1|	7

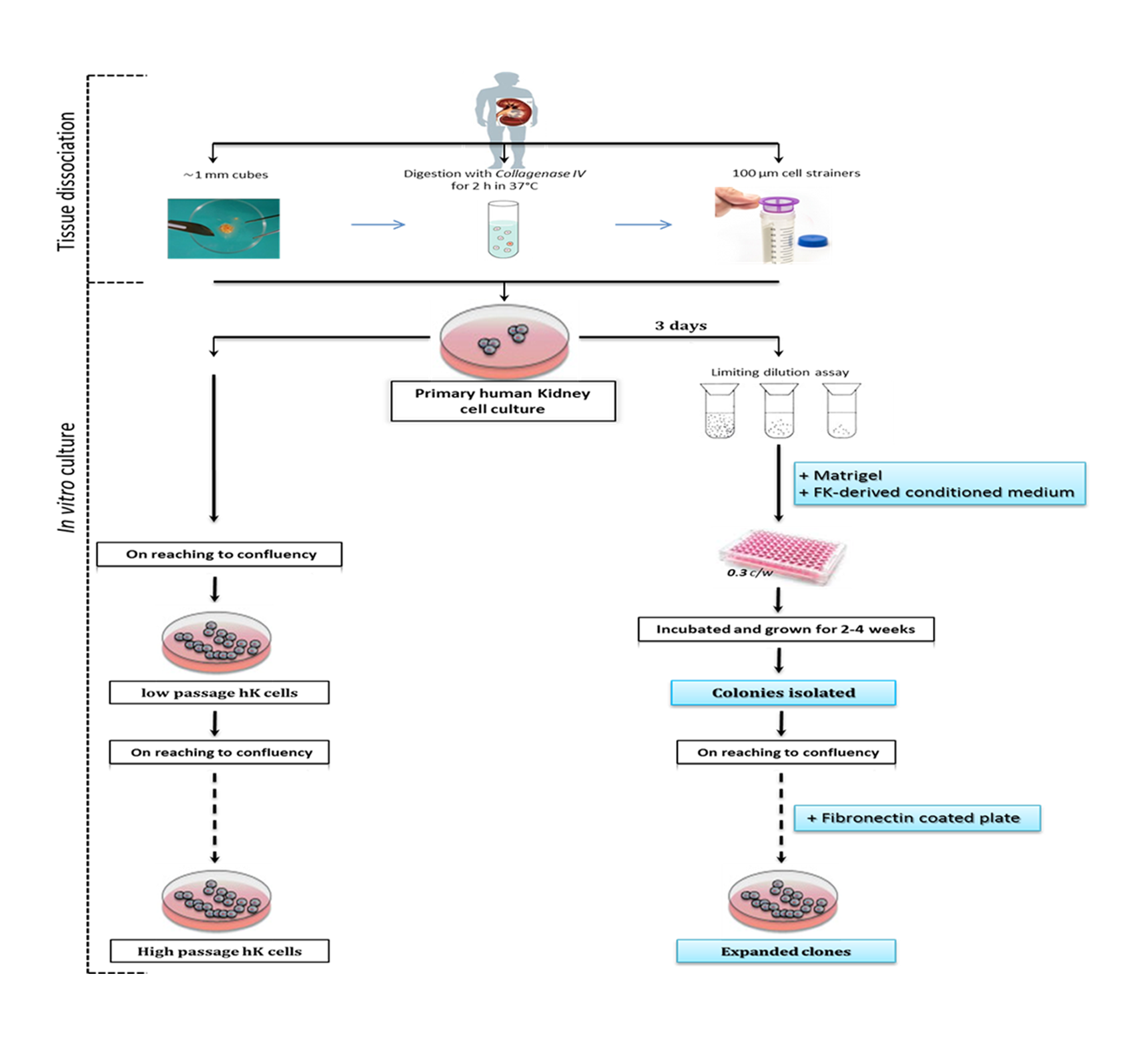

#### Supplemental figures and legends

##### Figure S1| Schematic representation of the method used for establishment of an efficient and reliable method for forming single cell clones from primary human

##### kidney. Flow diagram of the steps involved in adult kidney clones isolation and expansion.

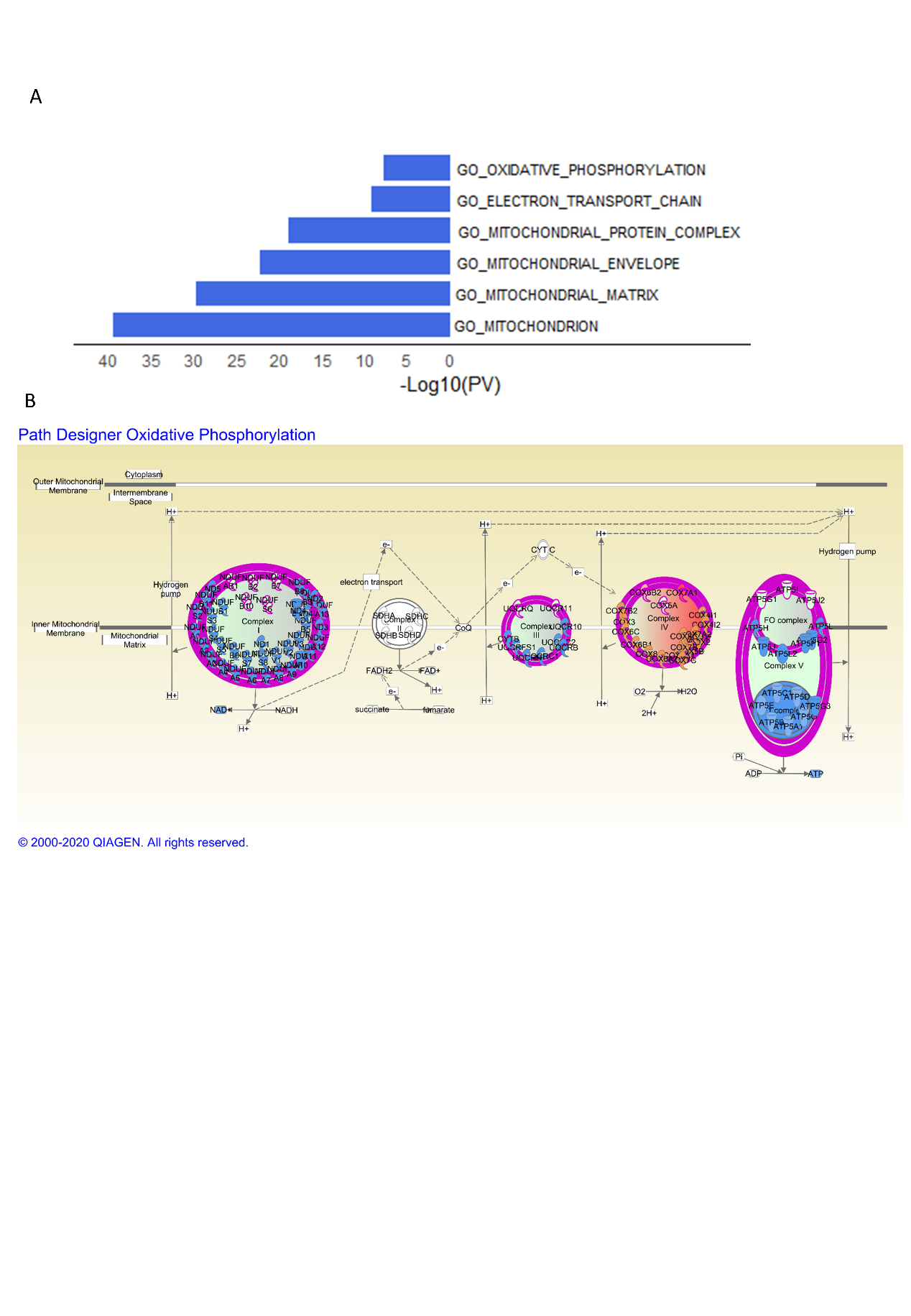

##### Figure S2| (A) Gene set enrichment analysis of differentially expressed statistics comparing ELC to FLC reveal an enrichment of mitochondrial-related GO terms in genes down-regulated in ELC (blue) and an enrichment of GO-terms related to development and specification processes in genes up-regulated in ELC (red); (B) Schematic representation of changes in mitochondrial related genes in FLC and ELC showing most complexes to be highly expressed by FLC. Annotations: Abbreviations: ELC- Epithelial-like clones, FLC- Fibroblast-like clones.

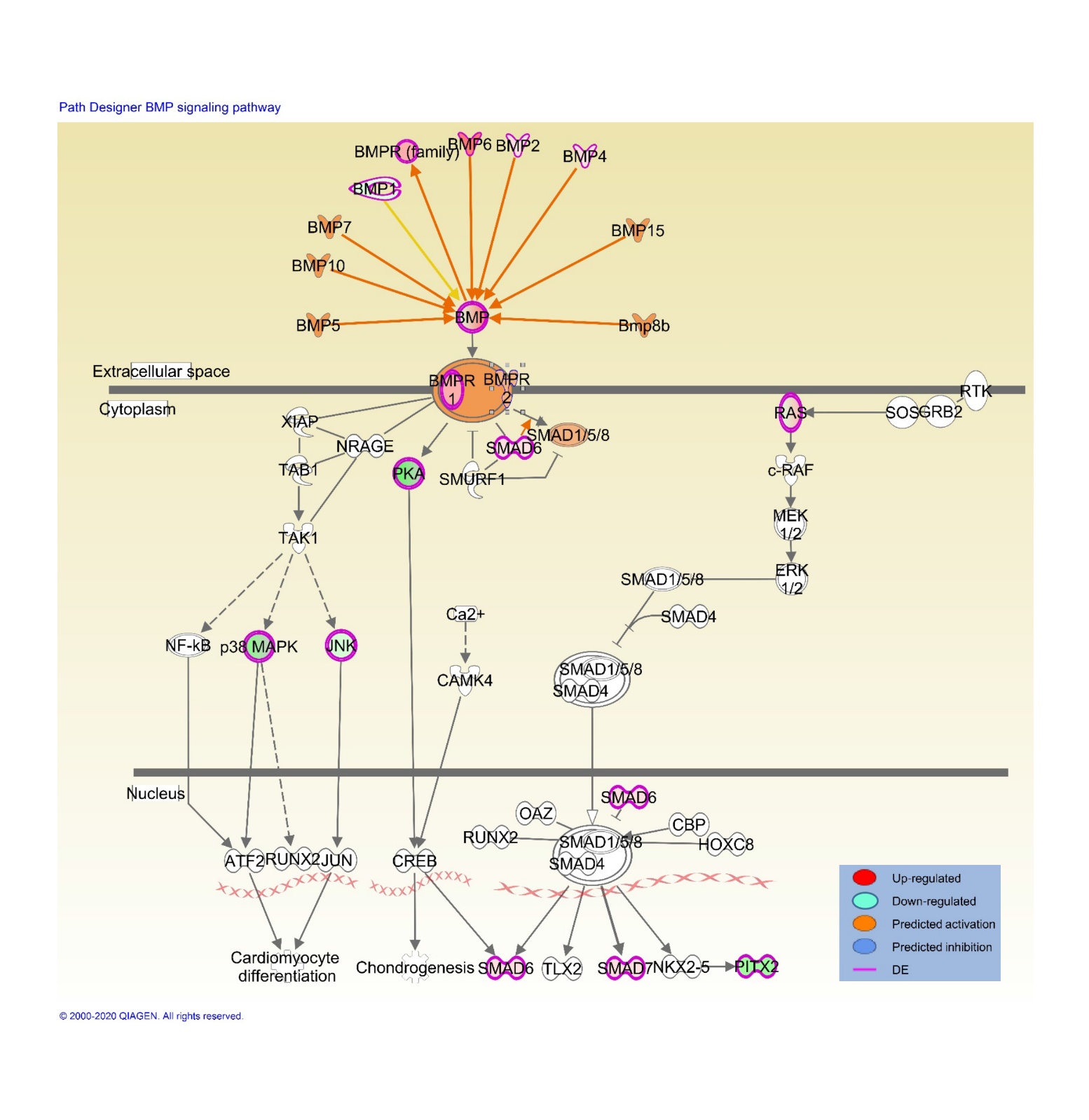

##### Figure S3| Schematic representation of changes in BMP/SMAD signal transduction pathway genes, showing many of its members to be highly expressed in ELC. Abbreviations: ELC-Epithelial-like clones

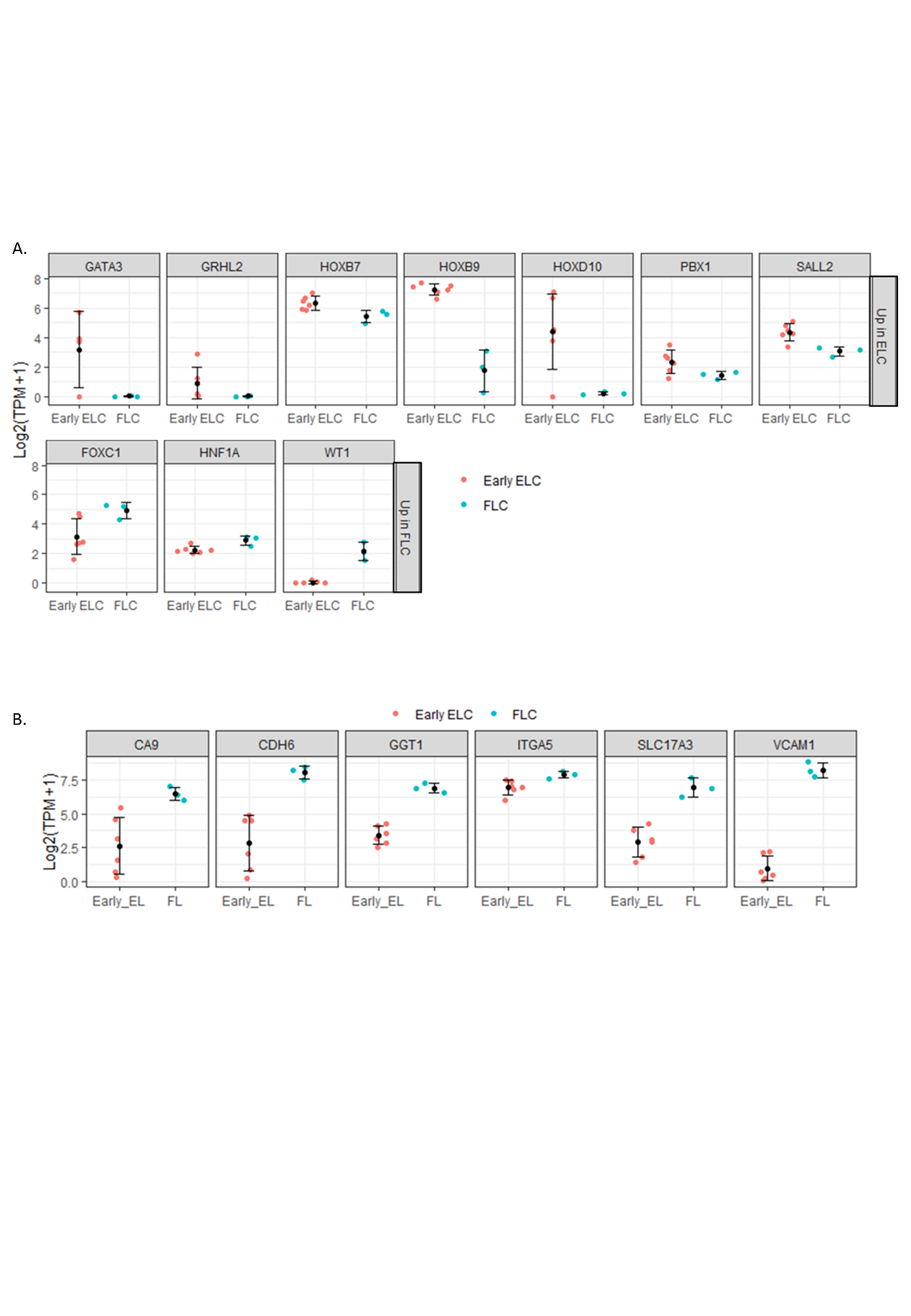

##### Figure S4| (A) Expression levels of nephrogenesis transcription factors differentially expressed between Early EL clones (red) and FL clones (turquoise). While genes upregulated in ELC are expressed in both the UB and the MM during renal development, genes upregulated in FLC are known to be expressed only in the MM; (B) Expression levels of Renal Cell Carcinoma (RCC) markers differentially expressed between Early EL clones (red) and FL clones (turquoise). Genes upregulated in FLC are known to be expressed in RCC.

#### Legend for Supplemental table

##### Table S1| Full RNA-Seq Data row counts used for the expression analysis of the different cultures (AK, ELC, FLC) generated from  Early or Late passages
